# TriLeukeVax: A CD80/IL-15/IL-15Rα Expressing Autologous AML Cell Vaccine Elicits Robust Anti-Leukemic Cytolytic Activity

**DOI:** 10.1101/2025.11.07.687000

**Authors:** Jacob Du, Uthpala N. Wijayaratna, Xinyue Wang, Jeffrey P. Fung, Belinda Huang, Farzin Farzaneh, Donald B. Kohn, Alexis J. Combes, Karin M. L. Gaensler

## Abstract

Acute myeloid leukemia (AML) is the most common acute leukemia in adults and is associated with poor outcomes due to frequent relapse after remission induction. While hematopoietic stem cell transplantation (HSCT) can improve survival, many individuals, especially older patients, are ineligible. Prior immunotherapies have not reliably induced effective anti-leukemic immunity and have been associated with severe and unpredictable toxicities. Thus, there is a need for safe and effective therapies that reduce relapse and increase overall survival (OS). We have developed a universally applicable, patient-specific, lentivirally engineered autologous AML cell vaccine, TriLeukeVax (TLV), designed to stimulate leukemia-specific cytolytic immune responses in AML patients in remission. To generate TLV, AML cells are engineered to express the highly synergistic combination of the co-stimulatory protein CD80 and the IL-15/IL-15-receptor alpha (IL-15Rα) heterodimer. Prior proof-of-concept (POC) studies demonstrated eradication of disease in >80% of leukemic mice with serial administration of TLV. In the current studies, TLV was generated from 59/60 cryopreserved, diagnostic bone marrow-derived patient AML samples. Ex vivo priming of post-remission patient T-cells by ex vivo co-culture with autologous TLV stimulated robust proliferative and cytotoxic responses. In secondary co-cultures, T-cells previously primed by initial co-culture with TLV, showed greater clonal expansion and leukemia-specific cytolytic activity towards de novo autologous AML blasts than did control, unprimed T-cells. The enhanced anti-leukemic activity of TLV-primed T-cells against de novo AML confirms the potential for vaccine administration to effectively target minimal residual disease (MRD) persisting after chemotherapy and reduce relapse.

**Key Points:**

- TriLeukeVax induces proliferation, activation, and effective anti-leukemic cytolytic responses in remission T-cells.
- Primed T-cells show polyclonal expansion and transcription profiles associated with proliferation, memory, and cytotoxicity.

Visual Abstract.
TriLeukeVax (TLV) workflow and mechanism of action.
Diagnostic AML patient bone marrow aspirates are collected, purified in Ficoll and cryopreserved. Samples are thawed and lentivirally transduced to produce TLV. Vaccination of AML patients in remission with TLV will stimulate the activation and expansion of leukemia-specific T-cells, effector memory cells, and NK cells by combining the co-stimulatory effects of CD80 with immune stimulation by the IL-15/IL-15Rα heterodimer expressed by the transduced AML cells, thereby targeting MRD and potentially increasing relapse-free survival in AML patients.

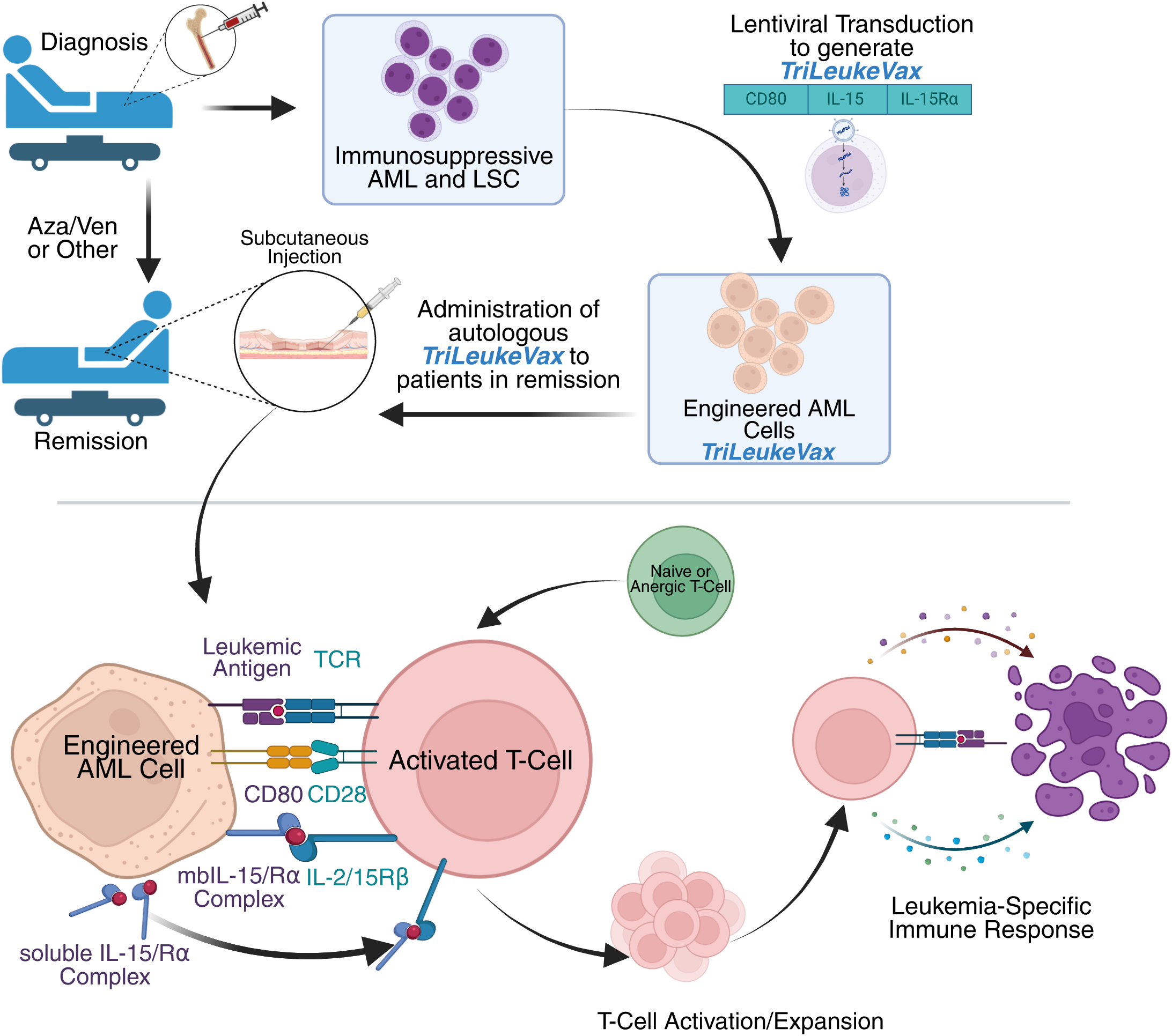

## 1. Introduction

AML is diagnosed in ∼23,000 adults/year in the US, with most occurring in patients > 60yo. Despite chemotherapy-induced remission rates of ∼60%, most patients relapse due to the persistence of quiescent leukemic stem cells (LSCs) and AML blasts.^1^ Allogeneic hematopoietic stem cell transplantation (HSCT) improves outcomes via donor Graft-vs-Leukemia effects targeting Minimal Residual Disease (MRD). However, even after HSCT, 30-50% of patients still relapse.^2^ Thus, more effective therapies are needed to target MRD and increase overall survival.

Current immunotherapies evaluated in AML patients have shown variable efficacy and unpredictable toxicities. Autologous leukemia cell vaccines may have significant advantages, as they have the potential to stimulate broad anti-leukemic immunity targeting multiple leukemia-associated and leukemia-specific antigens unique to the patient. This reduces the risk of immunological escape that occurs with therapies targeting single antigens. Compared to other immune-based therapies for AML, vaccines may also provide a safer option, particularly for older patients with poor performance status.^3–7^ Prior vaccines have shown promise and an excellent safety profile, but may have been limited by inefficient antigen presentation and poor induction of cytolytic activity.^8^ This is in part due to the down-regulation or absence of the critical co-stimulatory protein CD80 and other co-stimulatory ligands, combined with the elevated expression of co-inhibitory ligands on AML cells.

TLV is a universally applicable, patient-specific, lentivirally engineered autologous AML cell vaccine designed to stimulate leukemia-specific cytolytic immune responses in patients following remission induction. The vaccine is produced by lentiviral transduction of patient AML cells to express the synergistic combination of CD80 and IL-15/IL-15Rα, the naturally occurring dimeric cytokine that drives proliferation, activation, and persistence of cytotoxic T-cells and NK cells (Visual Abstract).^9^ This dual-stimulatory design directly addresses two central barriers to the success of immunotherapy in AML: (1) the lack of co-stimulatory signaling leading to T-cell anergy, and (2) the insufficient cytokine support and immune exhaustion of the host immune cells. By presenting the full repertoire of both private and public antigens on patients’ leukemia cells, TLV has the capacity to elicit broad polyclonal responses against a wide array of antigens without a priori knowledge of which antigens are most important.

In POC studies, a tricistronic lentiviral vector (LV) containing murine IL-15, IL-15Rα, and CD80 was used to transduce 32Dp210 murine leukemia cells to generate murine TLV (mTLV).^9–11^ Serial mTLV vaccination was curative in >80% of leukemic mice and showed efficacy in eradicating disease in a post-chemotherapy MRD model.^9^ Based on these results, the efficacy of this immune stimulatory approach was tested in ex vivo co-culture studies with TLV generated from patient-derived AML, and post-remission autologous T-cells. Priming of these T-cells with irradiated transduced autologous AML cells (TLV) induced strong polyclonal proliferative responses and cytolytic activity against unmodified AML cells in secondary co-culture assays.

## 2. Methods

### Lentivirus construction

The pMLV-hCD80-Furin-T2A-Opt-hIL-15-P2A-hIL-15Rα lentiviral vector (LV) was initially constructed by a homologous recombination procedure used to construct a previously published tricistronic murine IL-15-IL-15Rα-CD80-vector.^12^ Codon optimized human IL-15 and IL-15Rα sequences were provided by the Pavlakis laboratory (NCI).^13–16^ In prior published murine leukemia studies, this IL-15/IL-15Rα construct was incorporated to enable expression of membrane-bound and secreted IL-15/IL-15Rα dimers. The IL-15/IL-15Rα construct in the human tricistronic vector includes the human analog of this murine construct.^14^ The IL-15 cassette was linked by a P2A self-cleaving peptide sequence yielding hIL-15-P2A-hIL-15Rα. The TLV tricistronic LV was constructed by linking human CD80 to the IL-15-P2A-IL-15Rα via a furin cleavage site 5’ to the IL-15-IL-15Rα cassette, resulting in CD80-Furin-F2A-IL-15-P2A-IL-15Rα. The vector plasmid carries third-generation self-inactivating lentiviral sequences, containing a U3 deletion in the 3’ LTR and a SFFV-driven promoter in the 5’ LTR for production of LV RNA genome in transiently transfected producer cells. Pilot vector preparations were initially produced at Cincinnati Children’s Hospital Medical Center (CCHMC) Viral Vector Core, and subsequently optimized (Kohn Lab, UCLA). Engineering and GMP virus preps for IND enabling studies were produced at the Indiana University Vector Production Facility (IUVPF). A circular map of the human CD80/IL-15/IL-15Rα plasmid is shown (Supplementary Figure 1).

### AML patient sample collection, processing and cryopreservation

Bone marrow (BM), peripheral blood (PB), or leukapheresis samples were collected at diagnosis, and BM and PB were collected at remission in consented AML patients in accordance with the UCSF Institutional Review Board. Patient data derived from the electronic medical records was de-identified. BM mononuclear cells (BMMCs) were purified by Ficoll gradient centrifugation (Cytiva, USA) and cryopreserved in 90% heat-inactivated autologous plasma and 10% DMSO (Fisher, USA) or in CryoStor-10 (Biolife Solutions, USA) in a controlled rate freezer and stored in a liquid nitrogen freezer. Cryovials were thawed for 2 minutes in a 37 °C water bath, diluted in 5 mL pre-warmed AML thawing medium (RPMI 1640 supplemented with 10% human serum albumin (HSA), 10 μg/mL Pulmozyme, and 20 U/mL heparin), and incubated at 37°C, 5%CO_2_ for 30–60 minutes, followed by wash and cell culture.

### AML cell culture

The human AML cell line, U937 (ATCC, USA) and LV-transduced U937 (U937-TLV) cells were cultured in RPMI-1640 (Gibco, USA) with 10% fetal bovine serum (FBS, VWR, USA) and 100 U/mL Penicillin-Streptomycin (Gibco, USA). U937-TLV cells were prepared by lentiviral transduction of U937 cells, purified by fluorescence-activated cell sorting (FACS) (described below). Patient PBMCs and BMMCs were cultured in “AML Culture Media” (X-Vivo15 (Lonza, USA) with 50 ng/mL rhSCF, 50 ng/mL rhTPO, 50 ng/mL rhFlt3L, and 20 ng/mL rhIL-3 (Peprotech, USA)).

### Lentiviral transduction of patient AML cells

After resting in culture for 16-24 hours post-thaw, patient AML cells were transduced with the CD80/IL-15/IL-15Rα tricistronic LV (6E6 cells/mL, MOI=6.7 TU/cell) in AML Transduction Media (AML culture media containing 1mg/mL P338 Poloxamer (BASF Pharma) and 40.2E6 TU/mL lentivirus) and cultured at 37 °C for 16-20 hours. Cells were then diluted with fresh media, further incubated for 20-24 hours. Expression of IL-15, IL-15Rα, and CD80 was determined by flow cytometry.

### Irradiation of transduced primary AML cells

Transduced primary AML cells were irradiated (30Gy, RS 3400 X-Ray Blood Irradiator) at the UCSF Transfusion Service following standard operating procedures.

### Ex vivo co-culture assays

#### Primary co-culture assays

For ex vivo stimulation studies, cryopreserved post-remission patient PBMC/BMMCs were thawed and stained with anti-CD5 biotin (Biolegend, 300604) and purified by flow cytometry with streptavidin secondary antibodies, or by magnetic cell separation with anti-biotin microbeads (Miltenyi Biotech, 130-090-485). Purified T-cells were resuspended at 2E6 live cells/mL in AML Culture Media supplemented with IL-7 (10 U/mL) and rested overnight (37°C, 5%CO_2_). The following day (Day 1), transduced (designated “TLV”) and non-transduced (designated “AML”) AML cells were stained with CellTrace carboxyfluorescein succinimidyl ester (CFSE) (Invitrogen, C34554) and irradiated at 30Gy. Rested T-cells were stained with CellTrace Violet (CTV) (Invitrogen, C34557) to distinguish T-cells from AML cells in culture. CTV-stained T-cells were cultured alone, or with CFSE-stained, irradiated transduced (TLV) AML, or unmodified AML at 1:2 effector-to-target (E:T) ratio in IL-7-containing AML Culture Media. On Day 3, fresh media was added to each co-culture condition. On Day 5, cells were collected, stained, and T-cell proliferation and activation was assessed by flow cytometriy.

#### Resting of TLV-Primed T-cells

Following primary stimulation described above, TLV-primed T-cells were purified by FACS and dividing T-cells (Live/Dead^-^CFSE^-^CD3^+^CTV^dim^) collected. Dividing TLV-primed T-cells (2E5-2E6 cells/mL) were rested in AML culture media supplemented with IL-2 (30 U/mL), IL-7 (10 U/mL), and IL-15 (20 U/mL) for 7-10 days. Fresh media was added every 2-3 days.

#### Secondary co-culture assays

Cryopreserved AML patient PBMC/BMMCs collected at diagnosis were thawed and rested overnight. Non-primed T-cells were purified from thawed PBMC/BMMCs collected at remission and rested overnight (2E6 cells/mL in IL-7-containing AML Culture Media (37°C, 5%CO_2_)). On the following day, AML cells from diagnostic samples were cultured alone, with Non-primed T-cells, or with TLV-primed T-cells at 2:1 E:T ratio (2E6 live cells/mL in IL-7-containing AML Culture Media). Next day, cells were stained and analyzed for cytolytic activity by flow cytometry. Staining of primary cells with CTV or CFSE was performed according to the manufacturer’s protocols.

### Cell culture supernatant cytokine analysis

Cells were removed from culture and transferred into 96-well V-bottom plates, centrifuged at 500g for 6 minutes. Supernatants were transferred to 500 µL microfuge tubes, frozen, and stored at −80°C until shipment on dry ice. Cytokine analysis was performed using the Luminex™ 200 system (Luminex, Austin, TX, USA) by Eve Technologies Corp. (Calgary, Alberta). Fourteen markers were simultaneously measured in the samples (Eve Technologies’ Human High Sensitivity 14-Plex Discovery Assay®, MilliporeSigma, Burlington, Massachusetts, USA) according to the manufacturer’s protocol. Cytokines analyzed included GM-CSF, IFN-γ, IL-1β, IL-2, IL-4, IL-5, IL-6, IL-8, IL-10, IL-12p70, IL-13, IL-17A, IL-23, TNF-α, with assay sensitivities ranging from 0.11–3.25 pg/mL.

### Flow cytometry

Cells were incubated in FACS Buffer (2% FBS, 0.1% azide in phosphate buffered saline (PBS) for analysis, or 2% FBS, 2 mM EDTA in PBS for cell sorting) containing fluorophore-tagged panel of antibodies (Supplementary Table 3) and Fc-block for 30 minutes at 4°C. Samples that underwent intracellular staining or were not analyzed immediately were fixed using the eBiosciences FoxP3/Transcription Factor Staining Buffer Set (Invitrogen 00-5523-00). Stained cells were analyzed using the BD LSRFortessa™ or sorted using the BD FACSAria™. Data analysis was performed using FlowJo.

### Data and statistical analysis

Statistical analyses were performed using GraphPad Prism. Two-group comparisons utilized the unpaired student’s t-test. Three or more group comparisons utilized two-way or three-way ANOVA test followed by Tukey’s multiple comparison. Bar graphs represent sample mean +/- standard error (SE). All samples were analyzed in triplicate unless limited by primary patient-derived sample quantity.

Plasmid map figures were generated using SnapGene. Figure panels were created using BioRender.

### CITE-seq and T-cell sequencing

Samples were multiplexed into batches of 4-5 samples each. Diagnostic and remission samples from the same patient were pooled into separate batches. Cells were resuspended in PBS containing 0.04% bovine serum albumin (BSA). Cell counts and viability were determined using the Cellaca^TM^MX (Revvity). Equal number of cells were pooled from each sample to total 1E6 cells per pool. Pools were re-suspended in Cell Staining Buffer (Biolegend) and incubated with Fc Receptor Blocking Solution (Biolegend) at 20 mg/mL for 10 minutes on ice. Cells were then stained with a mixture of 140 TotalSeq-C oligo-tagged antibodies (Biolegend) for 30 minutes at 4°C. Stained cells were washed in PBS/1% BSA, filtered (40 µm cell-strainer), re-suspended in PBS/0.04% BSA, and loaded onto the Chromium Controller (10X Genomics) for Gel Beads-in-emulsion generation. cDNA libraries were generated using the Chromium Next GEM Single Cell 5’ v2 Library Kit for gene expression and the Chromium Single Cell 5’ Feature Barcode Library kit for antibody derived tags (10X Genomics). Enrichment of the V(D)J region was performed from cDNA, and additional libraries were constructed from these enrichments to generate TCR and BCR libraries. All libraries were subsequently sequenced on a NovaSeq S4 sequencer (Illumina) at the UCSF Center for Advanced Technology, supported by UCSF PBBR, RRP IMIA, and NIH 1S10OD028511-01 grants.

### Bulk RNA library preparation for demultiplexing CITE-seq and ATAC-seq libraries

Viably frozen PBMCs were thawed in RPMI1640 supplemented with 10% fetal calf serum at 37°C and cell counts and viabilities determined (Cellaca^TM^ MX). For each sample, 5E5 cells were aliquoted into RLT buffer (Qiagen). RNA was extracted from the lysed cells using Quick-RNA MagBead kit (Zymo Research) on the KingFisher automated extraction and purification system (ThermoFisher Scientific) according to manufacturer’s instructions. Extracted RNA was run on the fragment analyzer (Agilent) for quality assessment and quantification. cDNA libraries were generated using the Universal Plus mRNA-Seq with NuQuant kit (Tecan) following manufacturer’s instructions. The libraries were sequenced on HiSeq 4000 sequencer (Illumina).

### CITE-seq and TCR data analysis

CITE-seq data were pre-processed using CellRanger, DoubletFinder, and Seurat.^17–20^ Single cells were matched to patients with Freemuxlet, using patient-specific SNPs detected by bulk RNA-seq. To compensate for experimental bias and batch variation between sequenced samples, TLV-primed and non-primed T-cell datasets were integrated using the Harmony computation tool.^21^

## 3. Results

### 3.1. Successful generation of human tricistronic TLV lentiviral vector

To initiate studies with patient-derived AML and autologous PBMC, a lentiviral vector encoding human CD80, IL-15, and IL-15Rα was constructed and tested in AML cell lines and in primary patient AML samples. T2A and P2A cleavage sites separate the 3 genes. The Furin sequence preceding the 2A site enables efficient removal of the 2A residues.^22^ Expression of the codon optimized IL-15 and IL-15Rα sequences enables production of both membrane-bound and secreted IL-15/IL-15Rα heterodimers.^14,15^ The vector is designated pSFFV-hCD80-Furin-T2A-Opt-hIL-15-P2A-hIL15Rα (Figure 1A).

**Figure 1:**
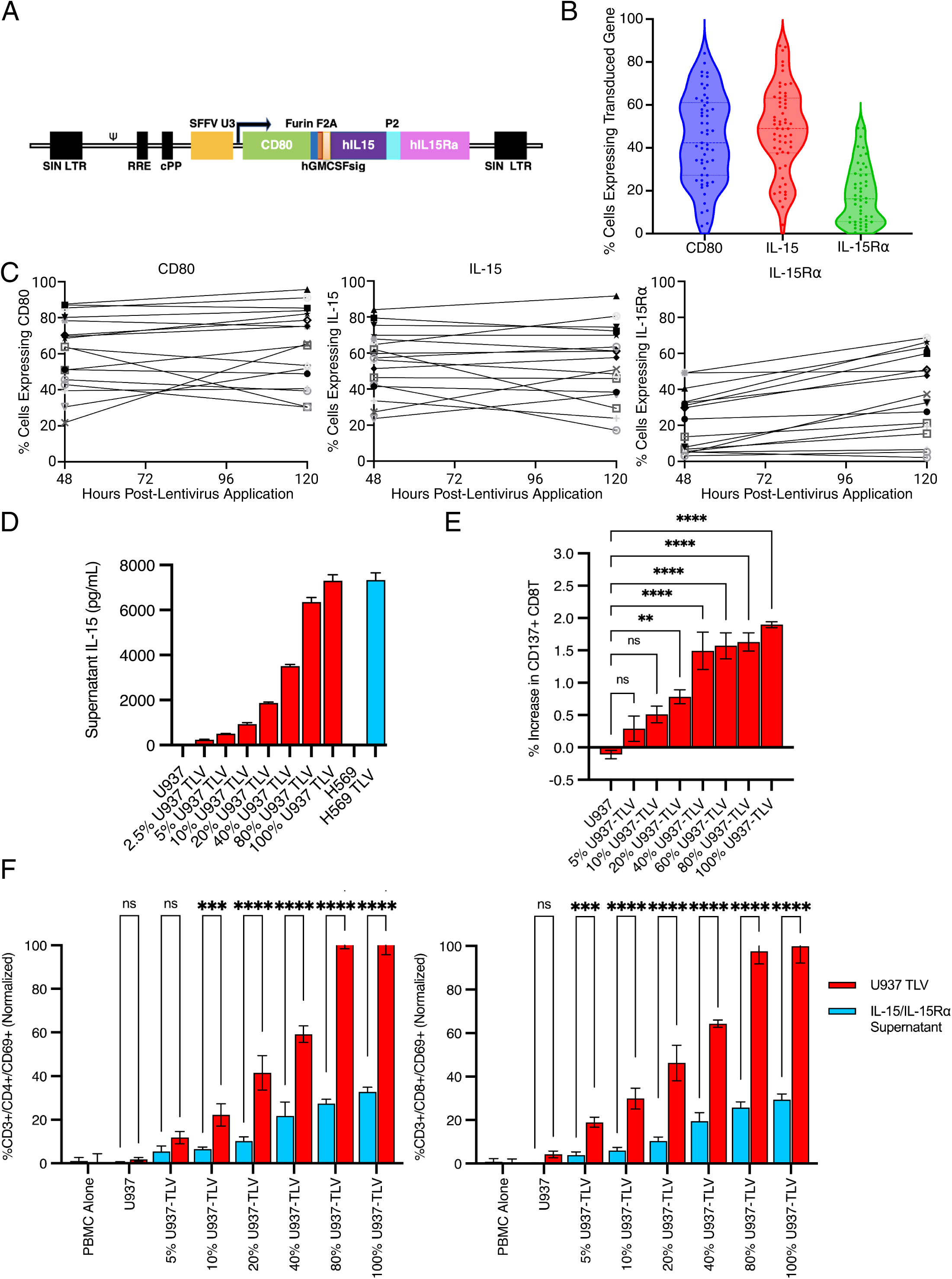
Characterization of TLV transgene expression and T-cell dose response. (A) Linear map of pSFFV-hCD80-Furin-T2A-Opt-hIL-15-P2A-hIL15Rα containing: (1) HIV-derived lentiviral vector of self-inactivating (SIN) LTR configuration, (2) ψ region vector genome packaging signal, (3) RRE (Rev Responsive Element) to enhance nuclear export of unspliced vector RNA, (4) cPPT (central PolyPurine Tract) to facilitate nuclear import of vector genomes, (5) SFFV U3 region promoter/enhancer, (6) Expression cassettes described above linked by 2A sequences for IL-15 and IL-15Rα, and Furin-2A for CD80 and IL-15. (B) Expression of transgenes CD80, IL-15, and IL-15Rα detected by flow cytometry after transduction of AML samples (36-44 hours after application of lentivirus) (n=60). (C) Expression of transgenes CD80, IL-15, and IL-15Rα detected by flow cytometry immediately after transduction of AML samples, or after 72 additional hours in culture in the absence of lentivirus. After transduction of AML samples, an aliquot of cells was taken for flow cytometry. Remaining cells were washed and cultured in AML media for an additional 72 hours before flow cytometry (n=17). (D) Levels of IL-15 secretion in transduced U937 cells and AML patient sample over 24 hours. Supernatants were collected and cryopreserved after 24 hours of culture using different mixtures of non-transduced U937 and U937-TLV (designated “TLV” on X axis in the graph to the right). Percentages reflect the frequency of U937-TLV in the cell mixtures. Patient H569 is a primary AML sample collected during therapeutic leukapheresis. No background IL-15 secretion was detected in non-transduced cells. (E) Dose/response assays with normal donor allogeneic PBMC stimulated with increasing fractions of transduced/non-transduced U937 cells for 5 days. Data for CD137 expression in CD8+ T-cells is depicted as % increase in expression compared to T-cells cultured alone. The U937 and U937-TLV cells were combined to create mixtures of 100%, 80%, 40%, 20%, 10%, and 5% transduced U937s. T-cells co-cultured with non-transduced U937s are labeled as “U937”. (F) Comparison of the level of PBMC activation as measured by CD69 expression in co-cultures of PBMC with irradiated U937-TLV versus culture with supernatants from U937-TLV cultures. Left panel shows CD69 expression in CD3+CD4+ T-cells; right panel depicts CD69 expression in CD3+CD8+ T-cells; red bars indicate co-culture with irradiated U937-TLV; blue bars indicate PBMC cultured in IL-15/IL-15Rα containing supernatants in the absence of U937 cells.

### 3.2. High level CD80/IL-15/IL-15R**α** expression achieved in AML

Pilot transduction of the human AML U937 cell line reliably directed high-level expression of all transgenes (Supplementary Figure 2A). By comparison, patient-derived AML samples were heterogeneous in terms of disease subtype, AML blast percentage, viability, and growth characteristics in culture (Supplementary Table 1 and Supplementary Figure 2B). As expected, the transduction efficiency in these samples varied, with 4.7%-84.1% CD80 and IL-15 gene expression (mean 48.2%, median 53.2%) (Figure 1B), as measured by flow cytometric analyses. Transduction efficiency was independent of AML cell subtype or variations in growth and viability in culture. The transduction efficiency of individual patient samples was reproducible with stable transgene expression for at least 120 hours in culture (Figure 1C).

To define the relationship between transduction efficiency and IL-15 secretion levels, IL-15 levels were quantified by ELISA in supernatants from cultures containing mixtures of transduced “U937-TLV” and non-transduced U937 cells. IL-15 levels were directly proportional to the fraction of transduced cells in the culture (Figure 1D). When IL-15 levels from a transduced patient-derived AML control sample (Patient #00569-TLV) were compared with supernatant IL-15 levels from 100% transduced U937-TLV, the levels were similar, demonstrating efficient secretion of IL-15 in primary AML-derived TLV.

### 3.3. Robust immune stimulation achieved with low level TLV transduction

Ex vivo dose/response studies were conducted to define the minimum level of AML transduction and transgene expression required to reliably stimulate T-cell activation. Allogeneic PBMC from a normal donor were co-cultured with titered mixtures of U937-TLV and non-transduced U937 at an E:T ratio of 1:1. After 5 days of co-culture, T-cell expression of CD137, a co-stimulatory receptor that is upregulated on activated antigen-specific T-cells, was quantified.^23^ Expression of CD137 by CD4+ T-cells was not analyzed because of the lack of MHC Class II expression in the U937 cell line. Control cultures of healthy allogeneic donor CD8+ T-cells with non-transduced U937 cells did not exhibit increased activation, defined by CD137 expression, compared to T-cells cultured alone. Dose-dependent increases in CD137 expression were observed in T cell co-cultured with up to ∼40% transduced U937 cells, with no further increases observed at higher transduction levels (Figure 1E). Stimulation with 10% U937-TLV yielded ∼27% of the maximal stimulation achieved with stimulation by 100% U937-TLV. Similar assessments were done to study the early activation marker CD69 after 24 hours of co-culture.^24^ Expression of CD69 is inducible by engagement of TCR, and by IL-15 and secreted factors from other activated T-cells. Quantifying CD69 expression enabled measurement of general activation of bystander T-cell populations, which may contribute to anti-tumor immunity via TCR-independent mechanisms.^25^ Co-cultures with 10% U937-TLV with CD4+ and CD8+ T-cells achieved 40% and 80% of the CD69 expression observed after stimulation with 100% U937-TLV (Supplementary Figure 2C). Thus, the potency of TLV is underscored by the reliable stimulation of immune responses with transduction levels as low as 10%.

### 3.4. Synergy of CD80/IL-15/IL-15R**α** co-expression in T-cell activation

In previous murine leukemia studies, co-expression of CD80 and IL-15/IL-15Rα showed high-level synergy in stimulation of leukemia-specific immune responses.^9^ To test the effects of combined CD80 and IL-15/IL-15Rα expression in stimulation of post-remission patient T-cells, co-cultures of patient-derived TLV with autologous CD4+ and CD8+ T-cells were performed. Levels of T-cell activation were compared between cultures stimulated with soluble IL-15/IL-15Rα alone (from U937-TLV culture supernatants) versus co-culture with defined mixtures of irradiated U937 and U937-TLV cells, where both CD80-mediated co-stimulation and IL-15 were present (Figure 1F). The frequency of activated CD4+ and CD8+ T-cells, defined by CD69 expression, was significantly higher after co-culture with TLV than after culture with supernatants containing IL-15/IL-15Rα from equivalent U937-TLV/non-transduced U937 ratios (Figure 1F). These results confirm the enhanced activity of combined CD80 and IL-15/IL-15Rα expression for immune stimulation of patient T-cells.

### 3.5. TLV stimulates proliferation and activation of autologous T-cells

Further co-culture studies were performed to test stimulation of autologous post-remission T-cells (Supplementary Figure 3) with autologous TLV generated by transduction of patient AML samples (Supplementary Table 2). TLV and autologous T-cells were stained with distinct fluorescent dyes prior to co-culture to distinguish the two populations. Irradiated (30Gy) TLV or unmodified AML cells were co-cultured with autologous T-cells at an E:T ratio of 1:2 for 5 days. Stimulation with TLV induced higher frequencies of T-cell activation, as measured by HLA-DR expression, a marker of late activation, and CD69 expression, than did co-cultures with unmodified autologous AML.^24^ This established the responsiveness of post-remission patient T-cells, even after prior administration of lymphodepleting chemotherapy (Figure 2A–B, Supplementary Figure 4). Multiplex cytokine analysis of a subset of patient samples revealed that TLV stimulation enhanced secretion of IFN-γ, IL-5, IL-13, IL-17A, and GM-CSF (Supplementary Figure 5A), consistent with broad immune activation. Co-culture with TLV also promoted greater proliferative responses in CD4+ and CD8+ T-cells, as measured by CTV dilution, when compared with unstimulated controls or co-cultures with unmodified AML cells (Figure 2C). Further, increased expression of CD56, a marker associated with effector T-cells with enhanced cytolytic potential, was detected in 5 of 7 patient CD8+ T-cell samples after TLV stimulation (Figure 2D).^26^ Similarly, increased expression of CD137 was observed in T-cells co-cultured with TLV (results achieving significance in 2/4 patient samples) (Supplementary Figure 5B). These results confirm the capacity of TLV to drive activation, proliferation, and effector differentiation of autologous AML patient T-cells, the future targets for immune stimulation by TLV administration in the clinical setting.

**Figure 2:**
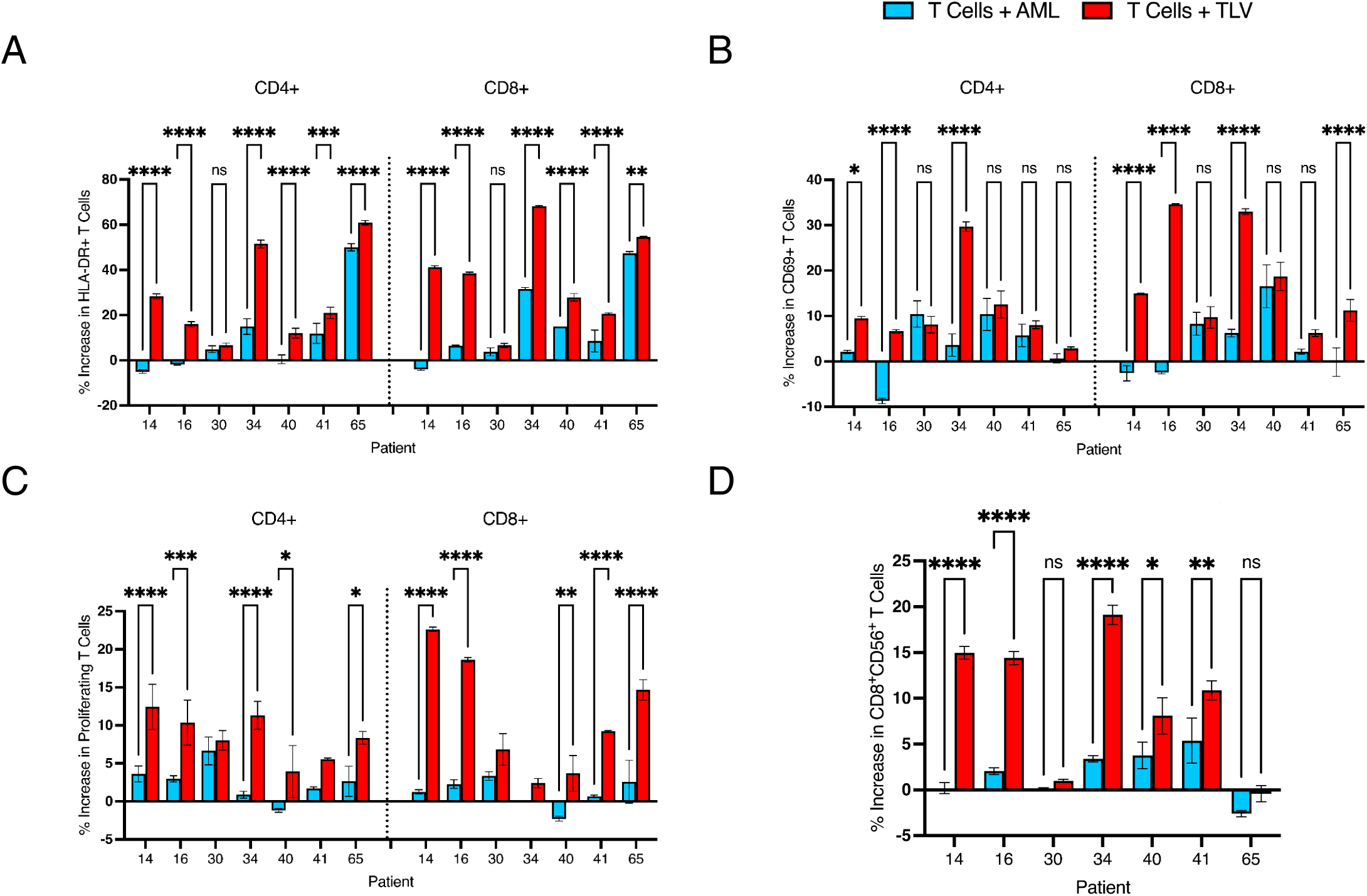
Assessment of T-cell activity after primary co-culture with TLV versus unmodified autologous AML shows enhanced activation and proliferation. Rested patient T-cells were stained with CellTrace Violet (CTV) and co-cultured for 5 days with irradiated, carboxyfluorescein succinimidyl ester (CFSE)-stained AML or TLV at an effector-to-target (E:T) ratio of 1:2. After the 5 day co-culture, cells were analyzed by flow cytometry. For analysis of primary co-culture samples, the % of dividing CD4+ or CD8+ T-cells was defined as the % of Live CTV+CD3+ cells with diluted CTV. The % increase shown in the figure was calculated by subtracting the average % dividing cells in T-cells cultured alone (background growth control) from those that were co-cultured. (A) and (B) Increased T-cell activation as measured by HLA-DR (A) and CD69 (B) after primary co-culture with TLV versus unmodified autologous AML. (C) Increased T-cell proliferation after primary co-culture with TLV versus unmodified autologous AML. The frequency of proliferating cells (CTV^dim^) was quantified in CD4+ or CD8+ T-cells (Live CFSE^-^CTV^+^) cultured alone or co-cultured with autologous AML or TLV. The increase in proliferation is calculated as % dividing T-cells co-cultured with AML or TLV, minus the %dividing T-cells in the absence of target. (D) Increased CD56 expression in CD8+ T-cells after primary co-culture with TLV versus unmodified autologous AML.

### 3.6. TLV induces AML-specific cytotoxicity and clonal T-cell expansion

Local TLV-mediated immune activation is intended to stimulate systemic anti-leukemic responses by clonal expansion of TLV-primed activated T-cells. To test the cytolytic activity of TLV-primed T-cells against residual autologous leukemia, secondary T-cell stimulation assays were designed by isolating responding proliferating T-cells from primary TLV co-cultures. Proliferating primed T-cells were purified by FACS and rested for 7-10 days to expand T-cell numbers and reduce background activation levels. Thereafter, the primed T cells were then tested in secondary co-culture with unmodified autologous AML (E:T 2:1 for 24 hours). The background cytolytic activity of non-primed T-cells was quantified by co-culture with de novo autologous AML. TLV-primed T-cells exhibited enhanced leukemia-specific cytolytic activity with decreased AML cell survival in secondary co-culture (Figure 3A, Supplementary Figure 6). To expand the analysis of TLV Primed T cell cytolytic activity, staining for intracellular cleaved Caspase 8 was employed. Caspase 8, an initiator caspase of apoptosis, is activated by engagement of T-cell death receptors such as Fas or granzyme B.^27^ Higher frequencies of intracellular cleaved-caspase 8 expression were detected in AML cells co-cultured with TLV-primed T-cells, establishing the robust induction of lytic T-cell responses against autologous AML with TLV stimulation (Figure 3B, Supplementary Figure 6).

**Figure 3:**
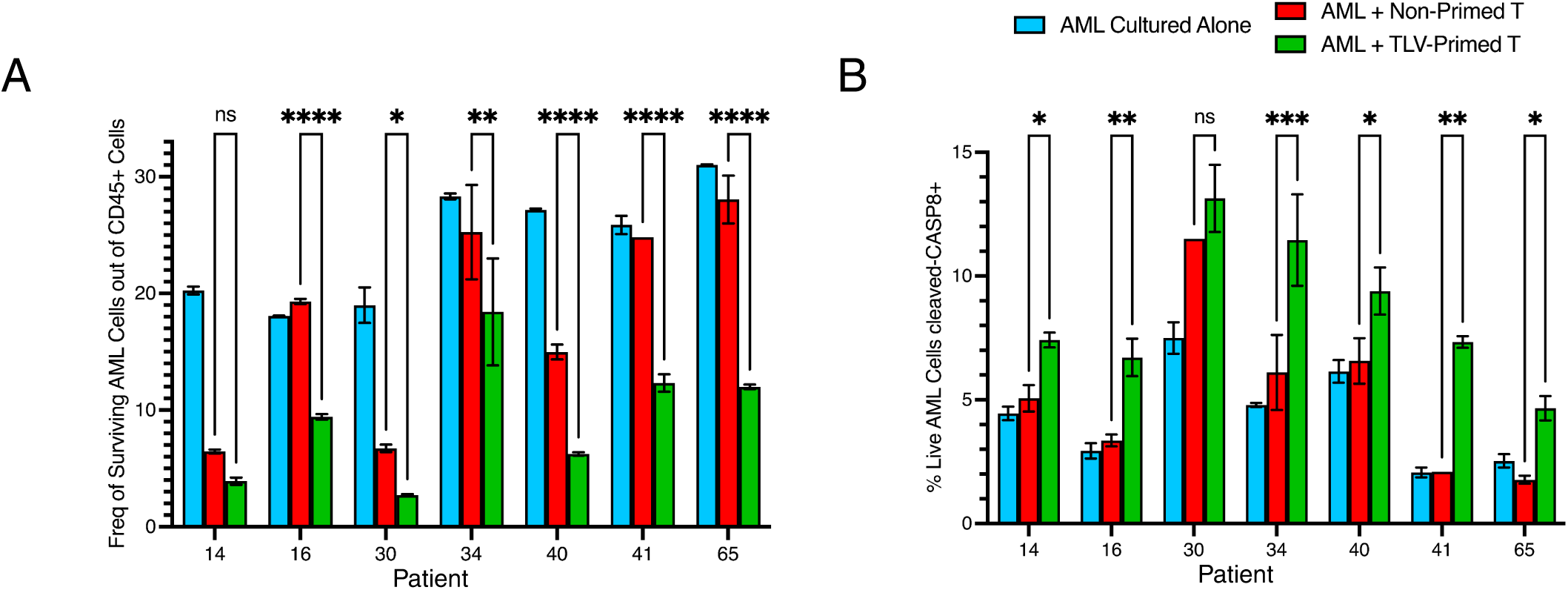
Priming with ex vivo TLV stimulation induces enhanced anti-leukemic cytolytic activity in patient T-cells. After purification of TLV-primed T-cells following primary co-culture, T-cells were rested for 7-10 days. Thereafter, T-cells were co-cultured with unmodified AML cells in secondary co-culture assays at an effector-to-target (E:T) ratio of 2:1 for 24 hours. As a control for baseline T-cell cytolytic activity, thawed non-primed T-cells were co-cultured with AML under identical conditions. For analysis of secondary co-culture samples, the % surviving AML cells was calculated by using the following formula: % Surviving AML = 100 * (# Live/Dead-CTV-CD45^dim^SSC^hi^ cells) / (# Live/Dead-CD45+ cells). In this analysis, CD45+ are leukocytes; CTV+ are primed or non-primed post-remission T-cells labeled with CellTraceViolet (CTV); and CD45^dim^ SSC^hi^ are AML or other myeloid cells. The formula above calculates the % of AML or other myeloid cells as a percentage (%) of live leukocytes, excluding primed/non-primed T-cells and other leukocytes originating from the BM sample present in co-culture. In Figure 2A, the % surviving AML in AML cells cultured alone (shown in blue) was normalized to the E:T ratios used in co-cultures. (A) Frequency of surviving AML cells after co-culture with Non-primed T-cells (Red) or TLV-primed T-cells (Green). The frequency of AML cells in peripheral blood or bone marrow samples cultured alone (Blue) was normalized to 33%, reflecting the 2:1 E:T ratio in co-cultures. (B) Percent increase in caspase-8 activity in AML cells was calculated by subtracting frequency of live AML cells with detected intracellular cleaved-Caspase 8 staining in co-cultures from those cultured alone.

In the clinical setting, the extent to which immune activation by TLV induces clonal expansion of leukemia-specific cytolytic T-cells is a critical question. To define transcriptional and TCR clonality changes in TLV-primed T-cells, CITE-seq studies were performed on purified, proliferating (n=7) and non-proliferating TLV-primed T-cell samples (n=2). Freshly thawed BM-derived T-cells (non-primed) from the same patients were sequenced in parallel. Unsupervised clustering of transcriptomes using the Leiden algorithm identified 16 transcriptional states, visualized by UMAP (Figure 4A) and annotated using common T-cell surface proteins (Supplementary Figure 7) and differentially expressed transcriptomes (Supplementary Figure 8).

**Figure 4:**
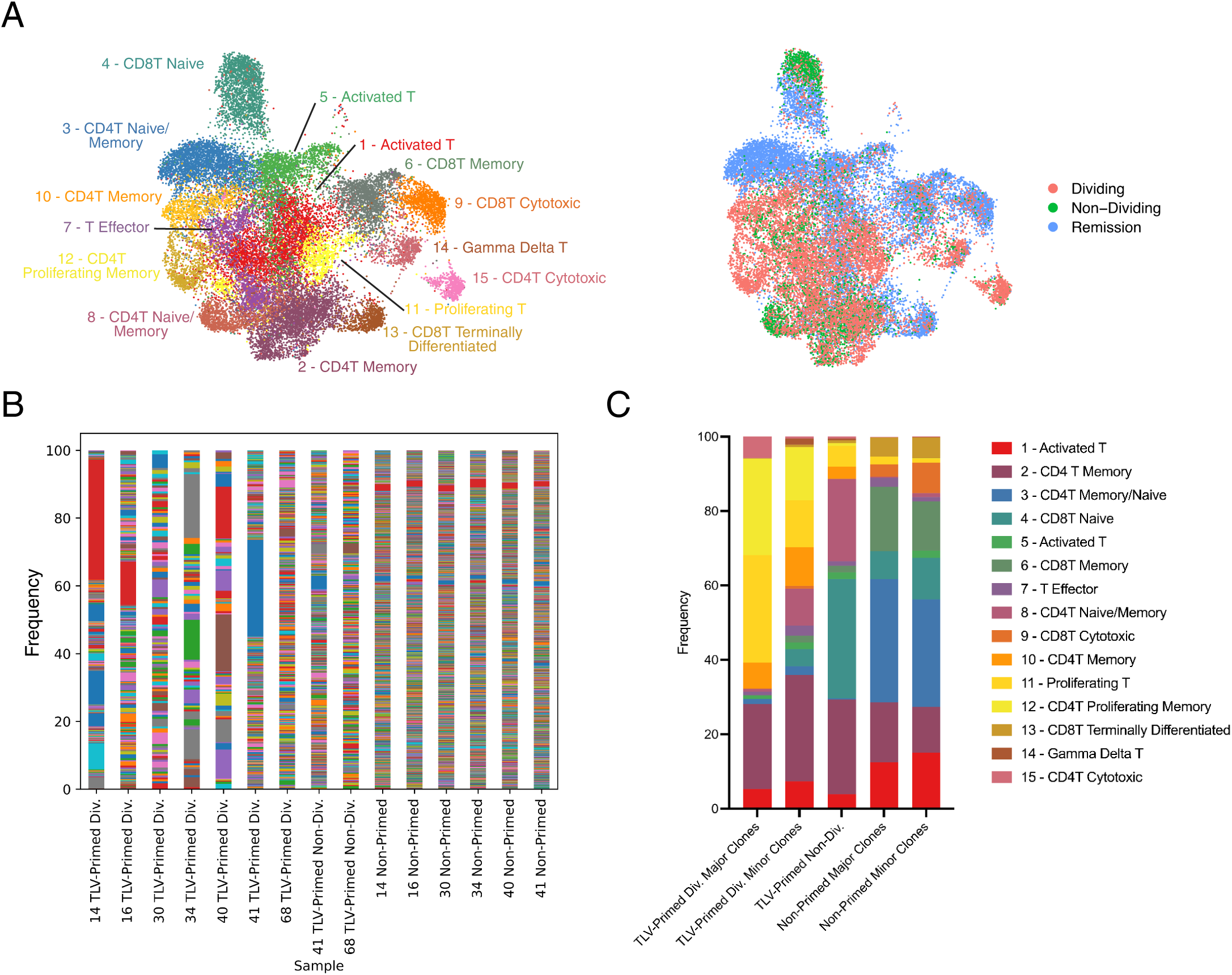
Transcriptome and clonotype analysis of TLV-primed T-cells show polyclonal expansion. (A) UMAP of non-primed and TLV-primed T-cells colored by T-cell subset (left) or sample type (right). After clustering cells with the Leiden algorithm (resolution of 0.8), the 16 identified cell clusters were annotated using differentially expressed transcriptomes and surface proteins. (B) Frequency of T-cell clones for T-cell samples sequenced. Each bar across the X axis represent a unique patient sample (identified by patient numbers 14, 16, 30, 34, 40, 41, 68 below the x axis). (C) Distribution of cell types of major and minor clones in dividing and non-dividing fractions of TLV-primed cells, and non-primed cells. Major clones consistent of clones representing >4% of the TLV-primed dividing T-cell samples (all other clones are considered minor clones). Clonotypes identified across samples belonging to the same patient were matched using TCR variable region CDR sequences.

TCR clones identified within each patient sample were further analyzed to compare T-cell clones among patients. Since samples for each patient were sequenced separately, identified clonotypes were matched between CITE-seq samples using variable region complementary determining region (CDR) sequences. Clonotype distributions within each sample were subsequently analyzed (Figure 4B). In 6 of 7 patients analyzed, one or more expanded clones (>4% of any given sample) were found in responding purified dividing TLV-primed T-cell samples, but were absent in the non-dividing TLV-primed and non-primed T-cell samples. These expanded clones were concentrated in three transcriptionally distinct clusters (#11, 12, and 15, Figure 4B). Cluster 11 is dominated by proliferation-related genes. Cluster 12 is characterized by a CD4 memory-associated phenotype (BATF3, GINS2, and NOP16) and by genes associated with proliferation. Cluster 15 is characterized by genes expressed in cytotoxic CD4 T-cells, including GNLY, PRF1, and IFNG. These transcriptional states were distinct from non-dividing TLV-primed T-cells, non-primed T-cells, and non-expanded clones within dividing TLV-primed T-cells (Figure 4C). The findings support the hypothesis that TLV stimulation induces polyclonal expansion of autologous T-cells exhibiting transcriptional profiles associated with proliferation, memory, and cytotoxicity.

## 4. Discussion

TLV was developed as a first-in-class, genetically engineered vaccine to address the need for effective therapies that reduce AML relapse. In these studies, successful lentiviral transduction of 59/60 independent primary BM-derived AML samples was achieved. Variable transduction efficiencies were observed between patients (4.7-84.1%, mean 48.16%, median 53.2%), likely attributable to the heterogeneity of the AML subtypes and patient population. However, transduction levels within individual sample replicates were reproducible. Importantly, in the setting of heterogeneous transduction efficiencies between samples, transduction of 10% of AML cells was established as sufficient to reliably activate autologous and allogeneic healthy donor PBMCs (Figures 1E-F; Supplementary Figure 2C). Modest differences (2-3-fold) in T-cell activation were observed using TLV samples with 10% to 100% transduction levels. IL-15 secretion levels were directly proportional to the transduced cell frequency. The high-level synergy of CD80 and IL-15/IL-15Rα co-expression in stimulating T-cell activation, previously observed with in vivo TLV vaccination in murine leukemia models, was largely recapitulated in ex vivo studies with patient samples. Outcomes of secondary co-culture assays support the hypothesis that TLV priming of T-cells enhances leukemia-specific T-cell cytolytic activity, modeling potential T-cell activation after serial vaccination in the clinical setting. Significant immune responses were observed even in T-cells harvested from older patients previously treated with lymphodepleting chemotherapy. Despite the expected heterogeneity of responses among the patient samples reflecting variations in age, AML subtypes, and other factors, TLV-mediated induction of anti-leukemic responses in autologous T-cells was quite effective, providing additional POC for the potential potency of TLV as a post-remission immunotherapy.

To begin defining transcriptional and clonal changes in T-cell populations after TLV stimulation, CITE-seq and TCR-seq were performed with TLV-primed and non-primed T-cells. Analyses showed polyclonal T-cell expansion in 6 of 7 independent TLV-primed dividing T-cell samples. This demonstrates that TLV can induce T-cell responses targeting multiple leukemia-associated antigens through presentation of both private and public antigens. Analysis of transcriptomes of the expanded clones revealed cell clusters/transcriptional states that were unique to, or enriched in, expanded clones in TLV-primed dividing samples. Expanded clones included a CD4+ T-cell cluster with a transcriptional state associated with cytotoxicity (high GZMB, PRF1, GNLY expression) that could be explained by MHC Class II (HLA-DR) expression in most AML cases.^28^

Recent studies have shown that engineered autologous tumor cell vaccines may have broader applications in other hematological malignancies, including multiple myeloma.^29,30^ Phase 1 clinical trials with other AML vaccine approaches have now established the feasibility of producing immunotherapeutics capable of stimulating leukemia-specific cytolytic activity in heavily pre-treated patients. As an autologous cell product, TLV presents the entire mutational landscape of patients’ AML, with the potential to elicit immune responses directed to multiple leukemia-specific antigens, many of which are unique to the patient. This reduces the risk of immunological escape mutants that occur with immunotherapies targeting single antigens, or even off-the-shelf allogeneic cell-based therapies that likely lack critical private antigens provided by an autologous drug. Therefore, TLV acts as a dual-function vaccine that enhances the breadth of induced immune responses by presenting both leukemia-associated antigens and immune-stimulatory signals intrinsic to the vaccine. Our studies support the feasibility of this approach and introduce the powerful cooperative effects provided by the combination of CD80 and IL-15/IL-15Ra expression for targeting residual disease and potentially preventing relapse.^29–35^ IND approval has recently been obtained for a Phase 1 study evaluating the feasibility, tolerability, and safety of TLV administration in AML patients achieving remission.

## Supporting information

Biorender Figure Publication License

## 5. Acknowledgments

The authors acknowledge Yimin Shi (University of California, San Francisco) for conducting prior murine studies and vector construction, Farzin Farzaneh (King’s College London, London, and Virocell Ltd. UK) for providing the CD80/IL-2 plasmid construct and for providing ongoing input and advice, George N. Pavlakis (National Cancer Institute) for providing codon optimized IL-15/IL-15Rα sequences, Jacqueline Tidball for extensive administrative support, and Donald B. Kohn (University of California, Los Angeles) for helpful discussions and collaboration in Pre-IND and IND enabling studies.

This work was supported by grants from the National Cancer Institute, National Institutes of Health (R21 CA177284-01A1) (K.M.L.G.); University of California, San Francisco–King’s Health Partners Faculty Fellowship Travel Grant (K.M.L.G.); University of California, San Francisco/Clinical and Translational Science Institute Pilot Project (K.M.L.G.); Leukemia and Lymphoma Society Translational Research Grant (6532-18) (K.M.L.G.); and California Institute for Regenerative Medicine (TRAN 1-11259 and CLIN 1-13985) (K.M.L.G.).

Sequencing was performed at the UCSF CAT, supported by UCSF PBBR, RRP IMIA, and NIH 1S10OD028511-01 grants.

Flow cytometry was performed at UCSF Parnassus Flow CoLab (RRID: SCR_018206), supported in part by Grant NIH P30 DK063720 and by the NIH S10 Instrument Grant S10 1S10OD021822-01.

## 6. Authorship

Contributions: J.D., U.W., X.W., J.F., B.H. performed experiments; J.D., U.W., X.W., and J.K. analyzed results and made the figures; J.D., X.W. U.W., A.C., and K.G. designed the research and wrote the paper.

Conflict-of-interest disclosure: K.G. has two approved patents (U.S. Patent No. US-12186342-B2 and U.S. Patent No. US-12453763-B2) and was listed as the inventor. The remaining authors declare no competing financial interests.

Correspondence: Karin M.L. Gaensler, Department of Medicine, University of California San Francisco School of Medicine, San Francisco, CA.

