## Supplemental Tables, Figures, and Figure Legends for "TriLeukeVax: A CD80/IL-15/IL-15Rα Expressing Autologous AML Cell Vaccine Elicits Robust Anti-Leukemic Cytolytic Activity"

Supplementary Table 1. Summary of AML patient disease subtype and AML blast % for samples used in these studies.

| Pt | AML Subtype | Clinical Flow Blast Frequency |
|---|---|---|
| 1 | AML consistent with therapy-related myeloid neoplasm | 70 |
| 2 | AML with monocytic differentiation | 1.1% myeloid blasts,56% monocytic, 9% imm. myeloid |
| 6 | Secondary AML converted from MDS / Recurrent/residual acute myeloid leukemia | 23% myeloid blasts, 20% monocytic, 9% imm. myeloid |
| 7 | Monocytic AML | 67 |
| 10 | AML | 38 |
| 11 | AML with mutated RUNX1, ASXL1 | 73 |
| 12 | AML with t(7;11) (FLT3 neg) | 96 |
| 13 | PM1+MDS/MPN overlap evolving to AML with extramedullary involvement in breasts c/w myeloid sarcoma | 0.1% myeloid blasts, 27% imm. myelomonocytic |
| 14 | Monocytic AML | 0.3% myeloid blasts, 58% monocytic |
| 16 | AML | 80 |
| 18 | AML (NPM1+) | 96 |
| 24 | AML | 82 |
| 25 | FLT3 ITD high, NPM1+ | Unknown |
| 26 | AML with monocytic diff | 92% myeloid blasts with monocytic diff. |
| 27 | Unknown | Unknown |
| 28 | Therapy related AML | 43% (additional 23% CD117+ CD34-) |
| 29 | Monocytic AML | 26% myeloid blasts, 47% monocytic |
| 30 | AML with monocytic diff | 58% myeloid blasts, 0.2% monocytic |
| 31 | monocytic NPM1+ AML | 24% myeloid blasts, 54% immature monocytic |
| 32 | Monocytic AML?? | 73 |
| 33 | AML | 33 |
| 34 | AML with monocytic diff | 38% myeloid blasts w/ monocytic diff |
| 35 | AML with monocytic diff | 23% myeloid blasts, 24% monocytic |
| 36 | Therapy related AML (Hx Breast Ca) +TP53, DNMT3A, WT1 | 89 |
| 38 | AML with monocytic diff | 1.6 |
| 39 | Secondary AML | 75 |
| 40 | AML (monocytic differentiation) | 46% blasts, 27% monocytic, 0.5% kappa restricted B cells |
| 41 | Monocytic AML with high FLT3 mutant allele fraction | 13.6% myeloid blasts, 59% monocytic |
| 44 | AML | 45 |
| 45 | AML FLT3ITD+ NPM1+ | 42 |
| 46 | AML | 77 |
| 47 | FLT3-ITD+ AML | PB-28% blasts |
| 48 | AML | 92 |
| 50 | FLT3+ AML | 61 |
| 51 | FLT3+ AML | 94 |

| | | |
|---|---|---|
| 54 | AML (monocytic differentiation) | 83% monocytic |
| 57 | Monocytic AML, FLT3 ITD+ | 86% monocytic |
| 61 | AML (MDS to AML) | 10 |
| 63 | AML tp53+ IDH2_ | 63 |
| 64 | NPM1+ monocytic AML | 30% myeloid blasts, 47% monocytic |
| 65 | FLT3-mutated AML | 85 |
| 66 | AML | 60 |
| 68 | inv 16 AML | 62 |
| 70 | AML ITD+ NPM1+ | 49 |
| 71 | AML with BCR ABL -> D15 CML with myeloid blast crisis | 24 |
| 74 | AML + monocytic differentiation | 8% myeloid blasts, 56% monocytic |
| 75 | AML with CEBPA+ | 73% |
| 76 | FLT3+ NPM1+ AML | 46 |
| 77 | AML | 24 (60% by morphology) |
| 79 | FLT3-ITD+, NPM1+ AML | 80 |
| 81 | AML | 62 |
| 83 | AML | 62 |
| 84 | TP53+ AML | 20 |
| 85 | AML | 25 |
| 86 | AML (monocytic differentiation) | 25% myeloid blasts, 11% monocytic |
| 87 | AML | 84 |
| 88 | AML (monocytic differentiation) | 47% myeloid blasts, 31% monocytic |
| 89 | AML with RUNX1::RUNX1T1 rearrangement | 32 |
| 90 | AML | 24 |

Pt indicates patient; AML indicates Acute Myeloid Leukemia; imm indicates immature

Supplementary Table 2. Summary of AML and remission T cell samples used in co-culture assays.

| Pt | Sex | Age | AML Subtype | Blast % | AML Source | Treatments (Dx→Rm) | T-Cell Source |
|---|---|---|---|---|---|---|---|
| 14 | M | 69 | NPM1+; Monocytic | 58% | Diagnosis BM | 7+3; entosplentinib | CR1 PB |
| 16 | F | 62 | TET2+, FLT3-ITD low | 55% | Diagnosis PB | 7+3; midostaurin | CR1 BM |
| 30 | M | 65 | MLL Rearr.; Monocytic | 58% | Diagnosis BM | 7+3 | CR1 PB |
| 34 | M | 65 | MLL+ Monocytic | 90% | Diagnosis BM | 7+3 | CR1 BM |
| 40 | M | 64 | Myelomonocytic | 80% | Diagnosis BM | Hydrea Decitabine/Venetoclax | CR1 BM |
| 41 | F | 47 | FLT3-ITD high; DNMTA3A+ | 73% | Diagnosis BM | 7+3; midostaurin | CR1 BM |
| 68 | F | 43 | CBFB::MYH11 rearrangement | 62% | Diagnosis BM | 7+3+ Gemtuzumab Ozogamicin (GO) | CR1 BM |
| 75 | M | 40 | RUNX1+ CEBPA+ | 73% | Diagnosis BM | 7+3 | CR1 BM |

Pt indicates patient; Dx indicates diagnosis; Rm indicates remission; PB indicates peripheral blood; BM indicates bone marrow; CR1 indicates 1$^{st}$ remission

Supplementary Table 3. Antibodies and dyes used for flow cytometry and ex vivo studies.

| Antigen/Marker | Clone | Manufacturer | Cat. No. | Conjugate/Fluor |
|---|---|---|---|---|
| Live/Dead | N/A | Invitrogen | L34987 | Scarlet 723 |
| CD5 | UCHT2 | BioLegend | 300604 | Biotin |
| CD45 | HI30 | Invitrogen | 47-0459-42 | APC-eF780 |
| CD80 | L307.4 | BD | 557226 | FITC |
| IL-15 | 34559 | R&D Systems | IC2471A | APC |
| IL-15Rα | JM7A4 | BioLegend | 330208 | PE |
| HLA-DR | L243 | BD | 564040 | BUV395 |
| CD3 | SK7 | BioLegend | 344812 | APC |
| CD69 | FN50 | Biolegend | 310914 | APC/Cy7 |
| CD56 | NCAM16.2 | BD | 564058 | BV786 |
| CD137 | 4B4-1 | Biolegend | 309804 | PE |
| Cleaved-CASP 8 | 18C8 | Cell Signaling Technologies | 9496L | None |
| Rabbit Igg H&L | N/A | Abcam | ab72465 | PE |
| CellTrace | N/A | Invitrogen | C34557 | Violet |
| CellTrace | N/A | Invitrogen | C34554 | CFSE |

Supplementary Figure 1

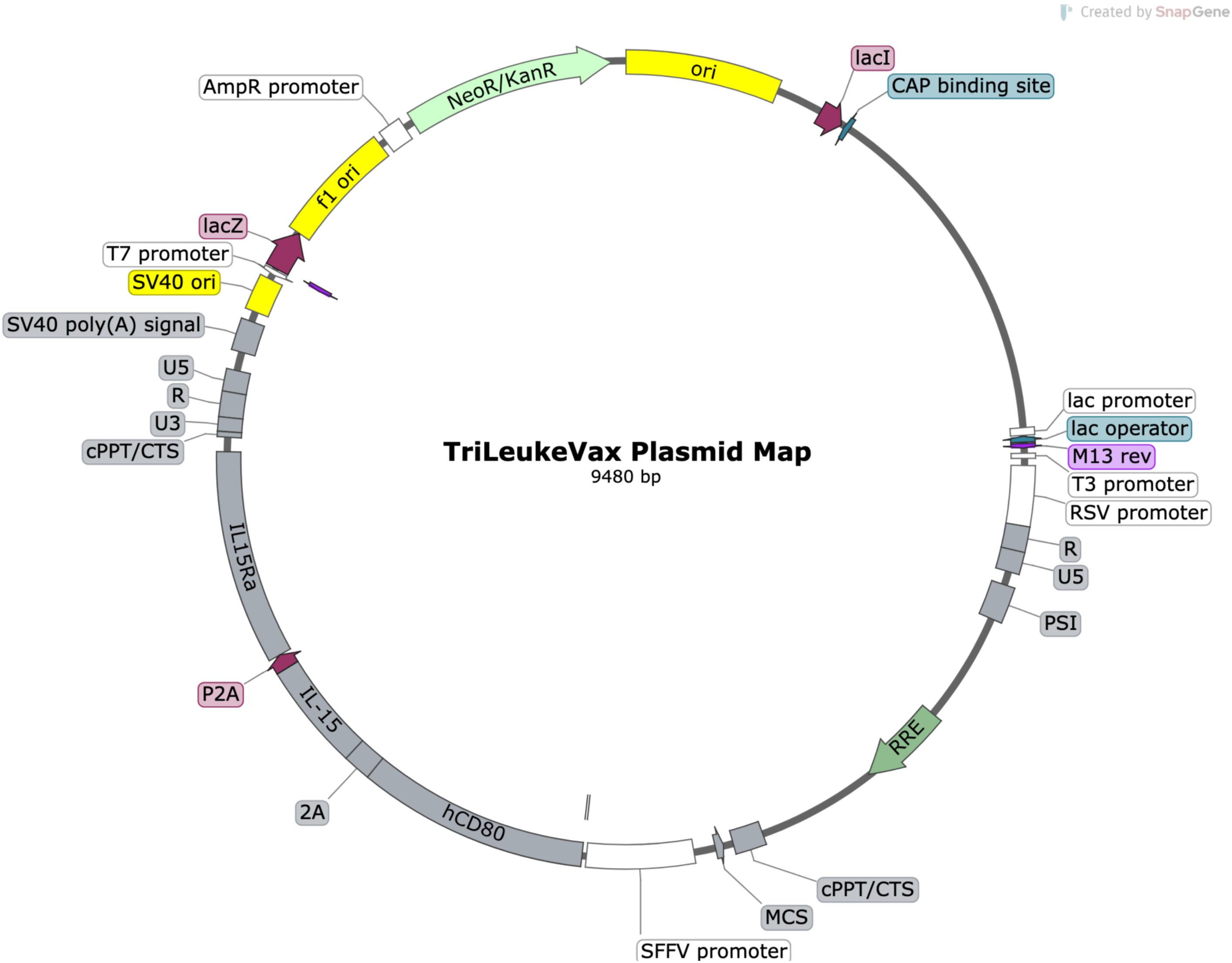

### Supplementary Figure 2

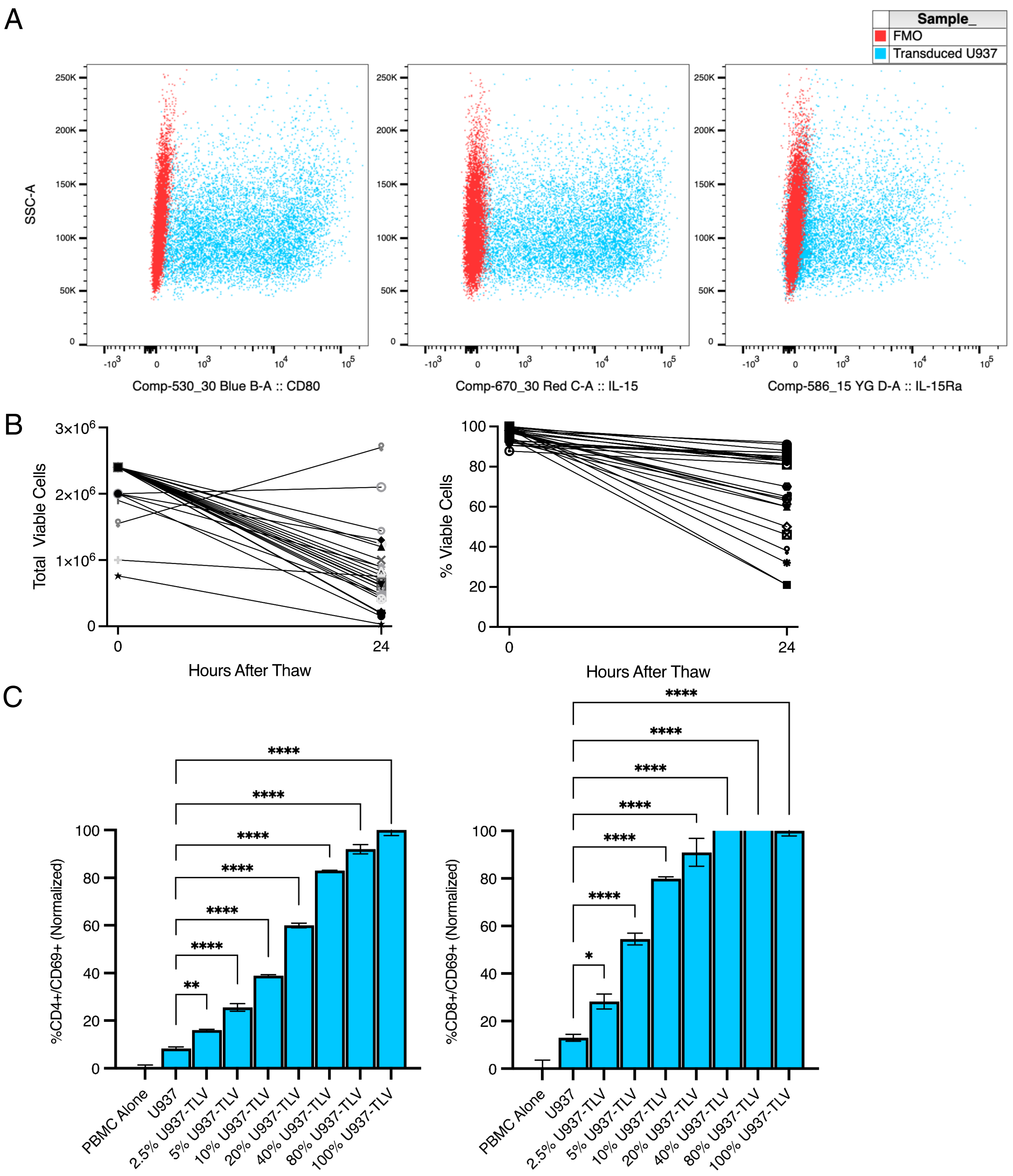

### Supplementary Figure 3

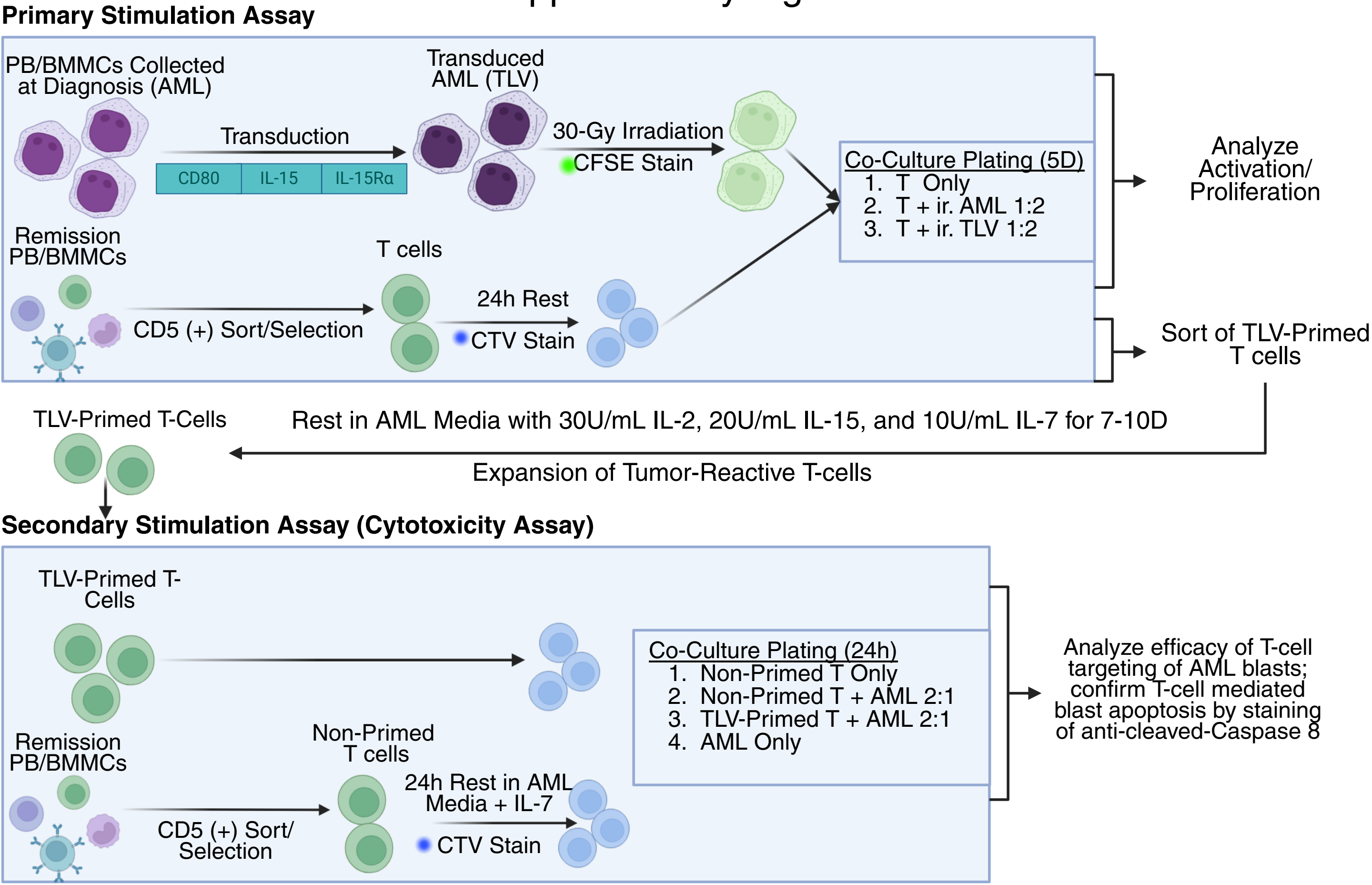

### Supplementary Figure 4

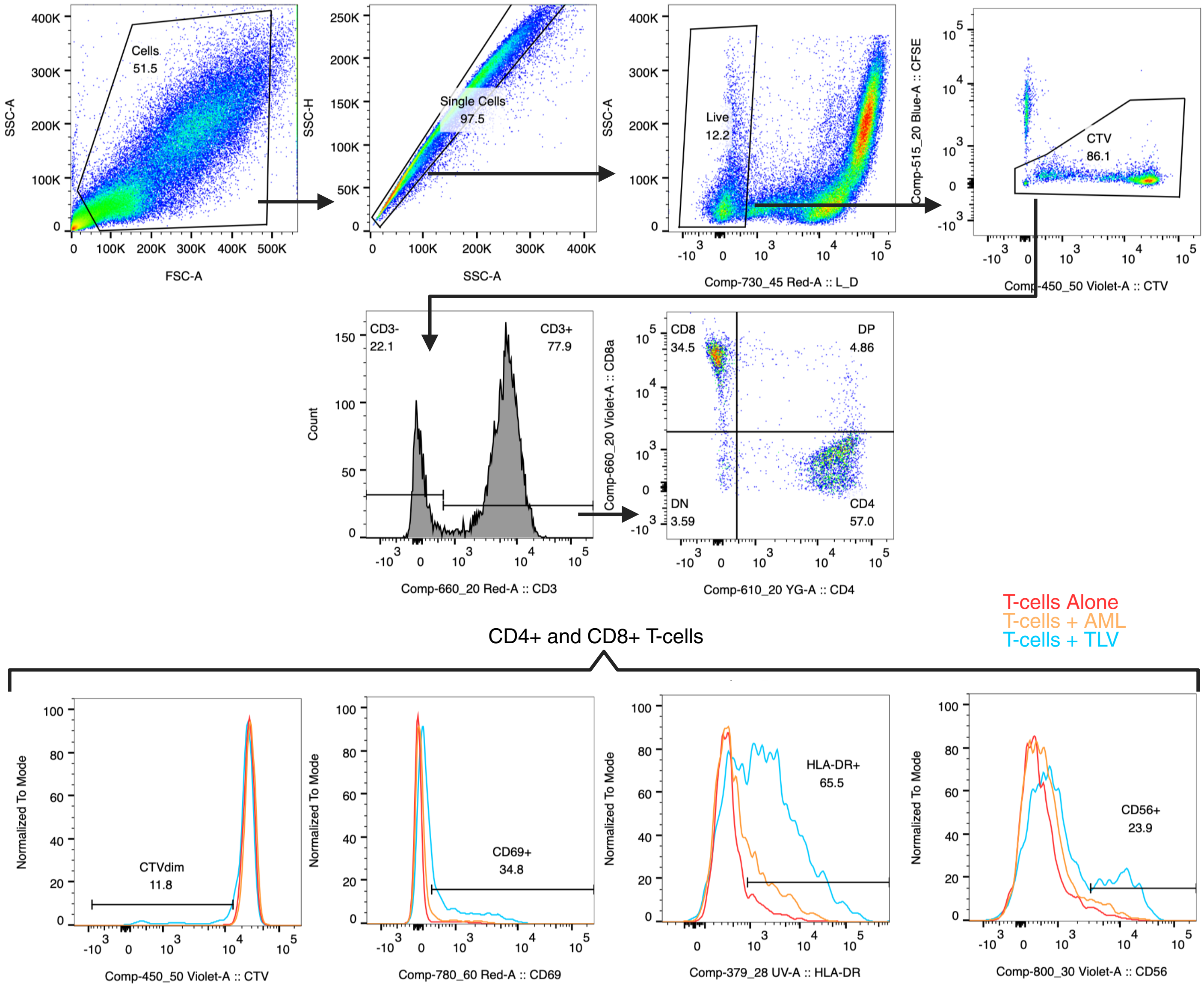

### Supplementary Figure 5

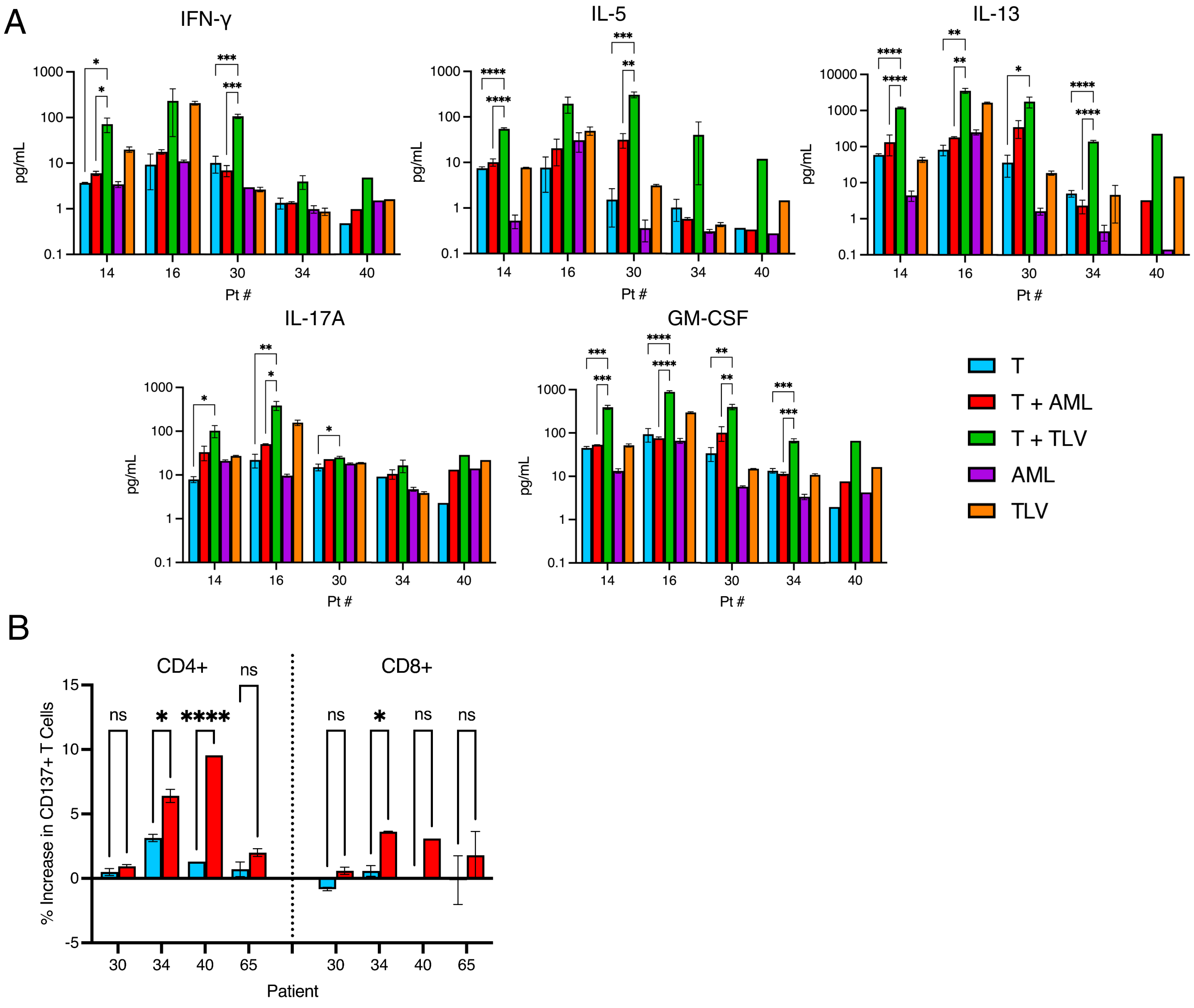

### Supplementary Figure 6

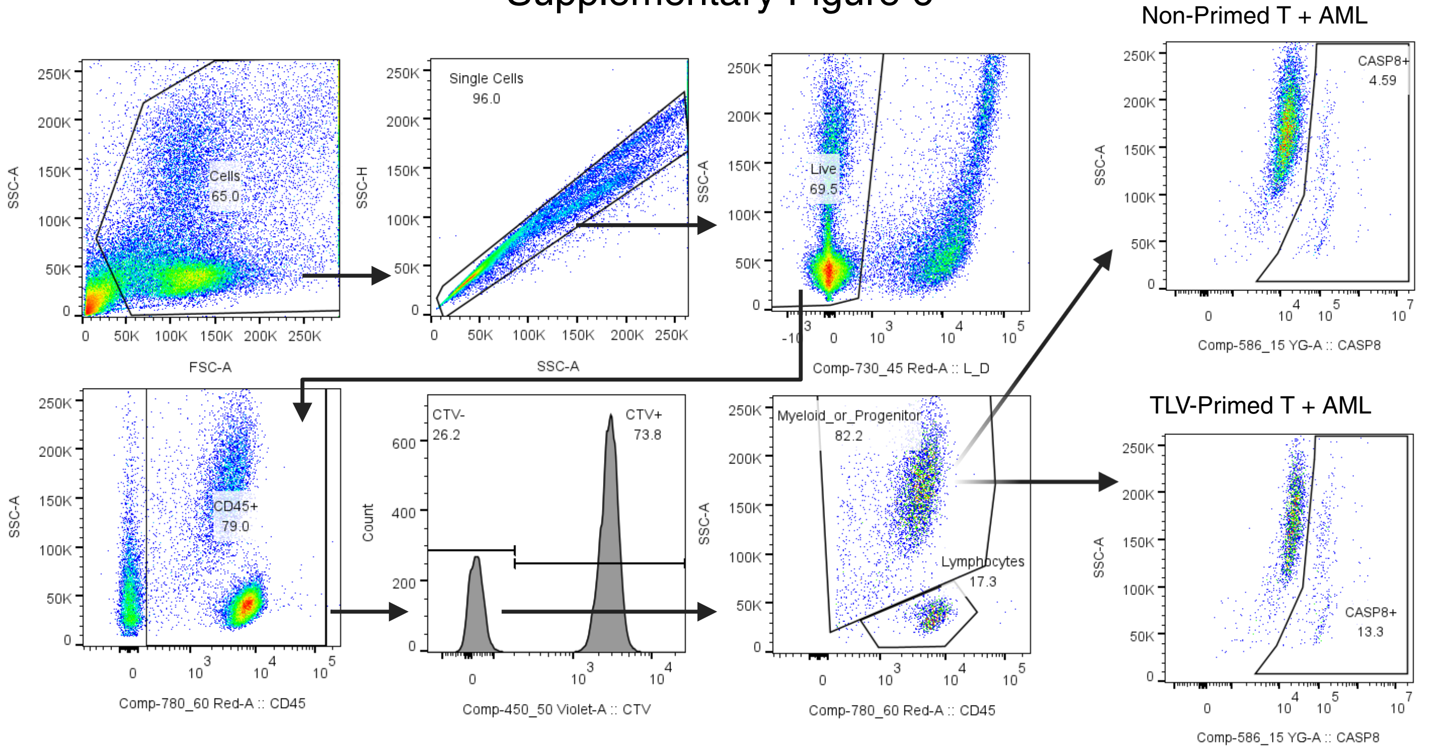

### Supplementary Figure 7

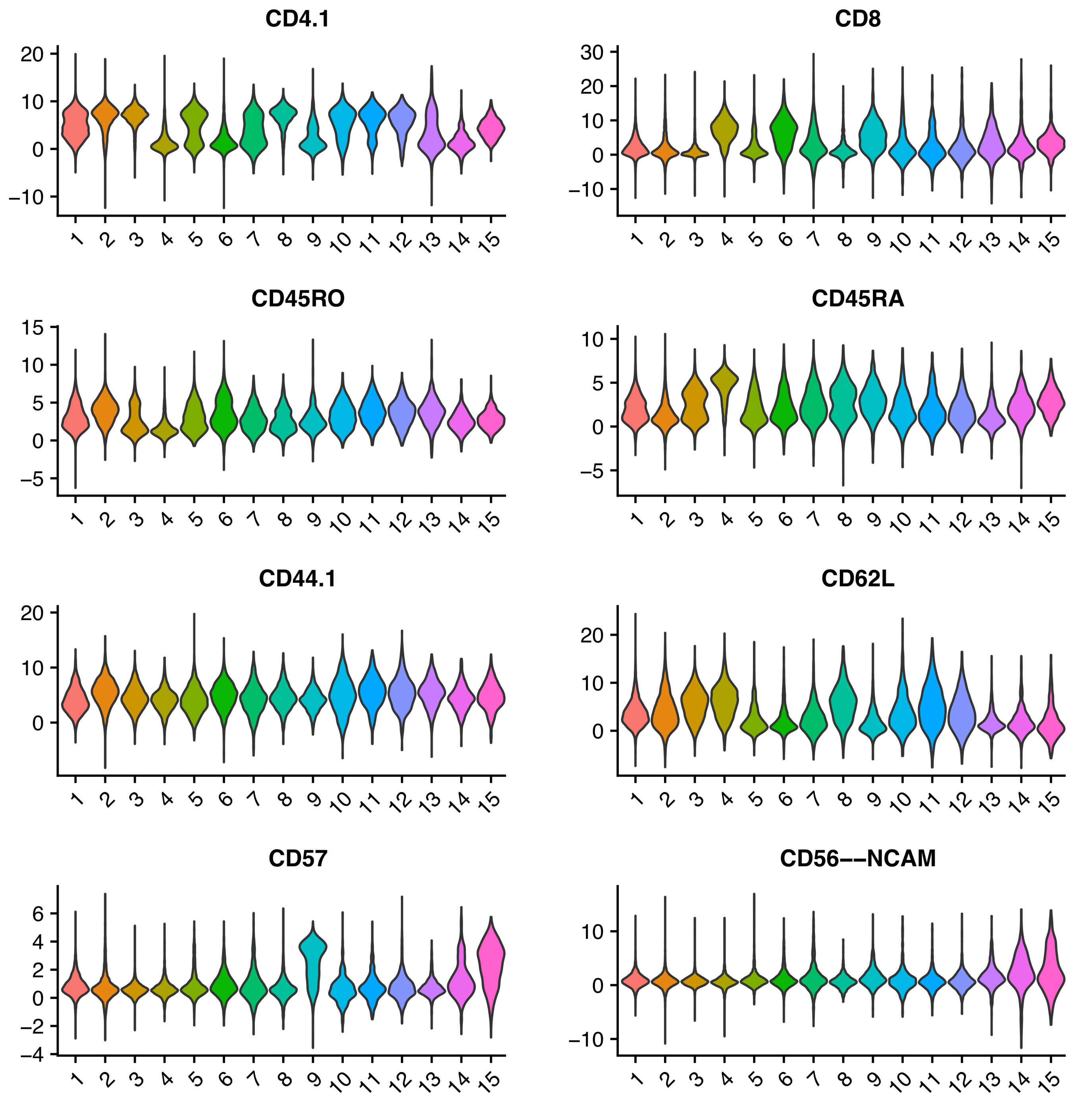

### Supplementary Figure 8

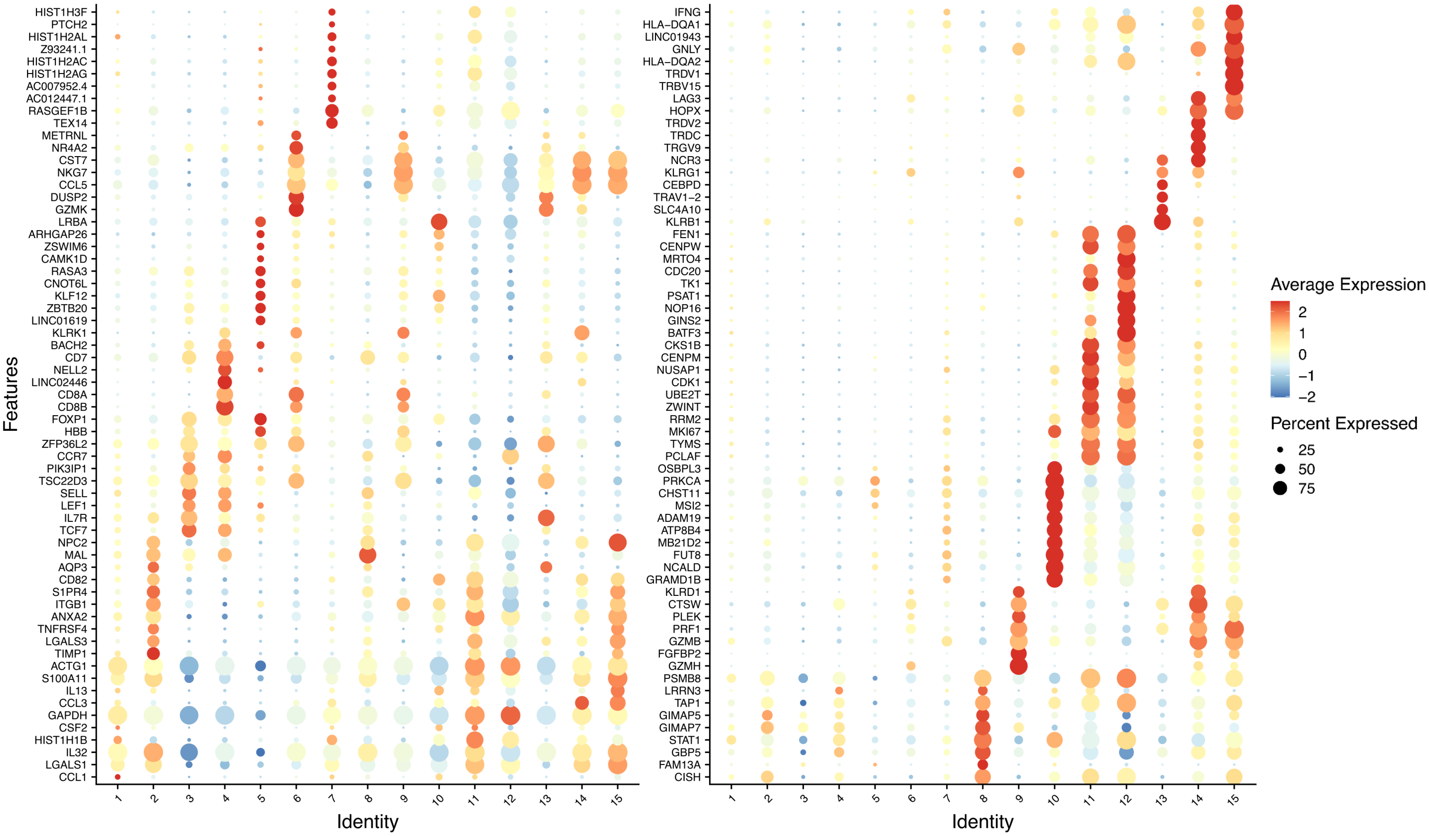

**Supplementary Figure 1: Plasmid map of TLV vector plasmid**. Annotated plasmid map representing the TLV vector plasmid generated in SnapGene Viewer.

**Supplementary Figure 2: AML cell and transgene expression characterization of TLV.** (A) Expression of transgenes in the human AML cell line U937 was analyzed by flow cytometry. U937 cells were transduced at an MOI of 6.7 TU/cell (as described in methods) and subsequently analyzed. Flow cytometry overlay plots show CD80, IL-15, and IL-15Rα expression on the X axis (left to right). Transduced U937 cells are shown in blue, and fluorescence-minus-one (FMO) controls are shown in red.

(B) Quantitation of bone marrow cell number (left) and loss of viability (right) over the first 24 hours after thaw (n = 27). Cell numbers and viability were counted using AO/PI fluorescent stains on the Nexcellom Cellometer Auto 2000.

(C) Dose/response assays with normal donor allogeneic PBMC stimulated with defined fractions of transduced and non-transduced U937 cell mixtures for 5 days. CD69 expression is shown in CD4+ (left) and CD8+ (right) T-cells. The U937 and U937-TLV cells were combined to create mixtures of 100%, 80%, 40%, 20%, 10%, and 5% transduced U937s. T-cells co-cultured with non-transduced U937 cells are labeled as "U937". PBMCs cultured in the absence of U937 or U937-TLV are labeled as "PBMC Alone".

**Supplementary Figure 3: Workflow diagram of primary and secondary stimulation co-culture experiments.** Upper panel depicts workflow of the primary stimulation co-culture assay while the lower panel depicts the workflow of the secondary stimulation co-culture assay (described in methods).

**Supplementary Figure 4: Gating trajectory used during analysis of co-cultures of T-cells with autologous TLV.** This figure shows the gating trajectory used for the analysis of T-cell proliferation and activation after primary co-culture with autologous TLV. T-cells are defined as Live/Dead- CellTrace CFSE- CD3+. CD4+ and CD8+ T-cells are then analyzed separately for CellTrace Violet (CTV) and other activation markers.

**Supplementary Figure 5: Primary stimulation assay cytokine analysis and assessment of CD137 expression.** (A) Quantitation of cytokine secretion during primary stimulation assays. Bone marrow or peripheral blood-derived T-cells were cultured alone or co-cultured with irradiated unmodified AML or TLV for 5 days at an E:T ratio of 1:2. As controls, AML and TLV were cultured alone in the same medium and at the same concentration. After co-culture, cytokines were analyzed as described in methods. Due to the 1:2 E:T ratio of co-cultures, contributions of cytokine secretions by irradiated AML or TLV cells quantified in T-cell co-cultures are 33% lower than the data shown for TLV or irradiated AML cultured alone.
(B) Increased T-cell antigen-specific activation as measured by increased expression of 4-1BB (CD137) after primary co-culture with TLV versus unmodified AML.

**Supplementary Figure 6: Gating trajectory used during analysis of primed T-cells with unmodified AML.** This figure shows the gating trajectory used for the analysis of TLV-Primed (and Non-Primed) T-cell cytolytic activity against autologous AML cells after secondary stimulation co-culture assays. The diagnostic peripheral blood/ bone marrow mononuclear cell fraction containing AML cells are defined as Live/Dead-CD45+CellTrace Violet (CTV)-. Myeloid or progenitor leukocytes (containing AML cells) are defined as $CD45^{dim}SSC^{high}$. Staining of cleaved-Caspase 8 is then analyzed in the myeloid or progenitor population.

**Supplementary Figure 7: Surface protein expression of identified cell clusters.** Violin plots of common T-cell surface proteins for identified cell clusters. Expression of these cell surface proteins (CD4, CD8, CD45RO, CD45RA, CD44, CD62L, CD57, and CD56) were used for annotation of identified cell cluster (depicted as #1-15 on the X axis).

**Supplementary Figure 8: Differentially expressed transcriptomes of identified cell clusters.** Dot plot of top 10 differentially expressed transcriptomes (DEG) of each identified cell cluster. After cell clustering, the DEGs of identified cell clusters were calculated based on % expression of genes within a cluster relative to all other cells. The dot plot shown was generated using the top 10 DEGs for each cluster (depicted as #1-15 on the X axis).
