## Supplementary material for "TriLeukeVax: A CD80/IL-15/IL-15Rα Expressing Autologous AML Cell Vaccine Elicits Robust Anti-Leukemic Cytolytic Activity": Biorender Figure Publication License: 2 - Publication License Nov-05-2025.pdf

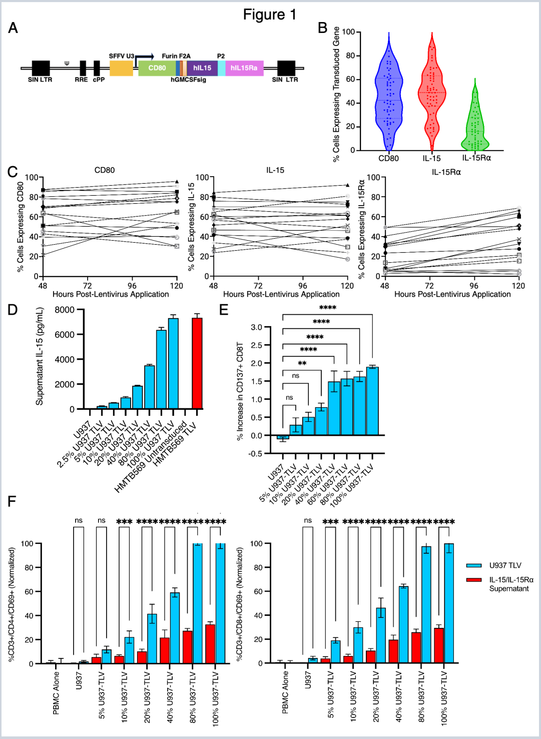


*For any questions regarding this document, or other questions about publishing with BioRender, please refer to our [BioRender Publication Guide], or contact BioRender Support at [].*
