## Supplementary material for "TriLeukeVax: A CD80/IL-15/IL-15Rα Expressing Autologous AML Cell Vaccine Elicits Robust Anti-Leukemic Cytolytic Activity": Biorender Figure Publication License: 2 - Publication License Nov-05-2025.pdf

49 Spadina Ave. Suite 200
Toronto ON M5V 2J1 Canada
www.biorender.com

### Confirmation of Publication and Licensing Rights - Open Access

**November 5th, 2025**

| | |
|---|---|
| ***Subscription Type:*** | *Individual - Academic* |
| ***Agreement number:*** | *IJ28YME3S6* |
| ***Publisher Name:*** | *Biorxiv* |
| ***Figure Title:*** | *Supplementary Figure 1-8* |
| ***Citation to Use:*** | *Created in BioRender. Du, J. (2026) https://BioRender.com/wa0h86e* |

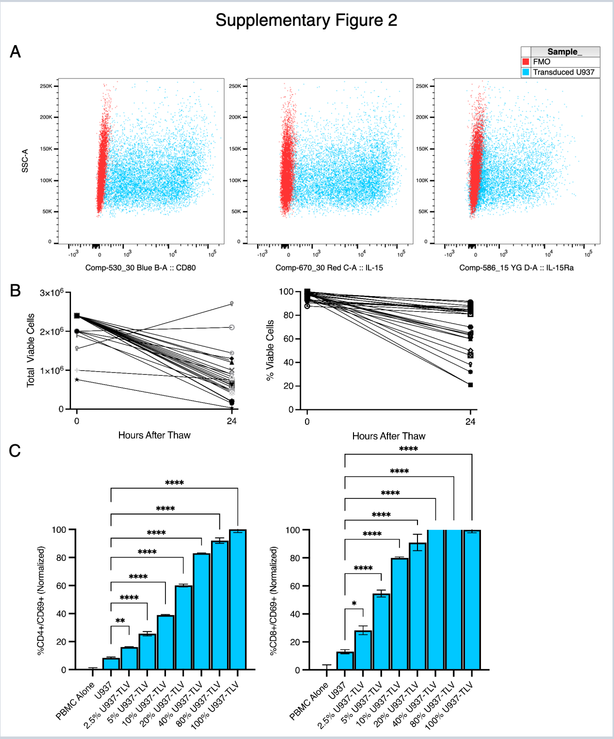


*For any questions regarding this document, or other questions about publishing with BioRender, please refer to our BioRender Publication Guide, or contact BioRender Support at.*
